# Proteins with multiple G protein-coupled receptor domains

**DOI:** 10.1101/2022.07.26.501653

**Authors:** Kilic Isildayancan, Amit Kessel, Ron Solan, Rachel Kolodny, Nir Ben-Tal

## Abstract

Currently known G protein-coupled receptors (GPCRs) have a single transmembrane domain. Many GPCRs form dimers that have two transmembrane domains (one per protein), and there are indications that this dimeric interaction is functionally meaningful. Here, based on sequence analysis and structure predictions, we report the existence of 57 proteins with two, three, or four GPCR domains within the same protein chain. We analyze the structures of these multi-GPCRs and show that almost all have DRY/NPxxY motifs, a strong indication of signaling activity. By homology, most of the multi-GPCRs that we identified are olfactory-related; a few are chemokine-related. Multi-GPCR candidates are found in various Chordata species including fish, camel, marmite, Chinese hamster, and new world monkeys. The discovery of receptors with multiple transmembrane domains suggests the possibility for signal regulation and amplification within an individual receptor, revealing another step in GPCR evolution and a new layer of complexity in signal transduction.

## Introduction

GPCRs, also known as G protein-linked receptors, are by far the most diverse and broadly distributed family of membrane-resident receptors [1]. The genes encoding GPCRs constitute 1-5% of vertebrate genomes [2-4], and there are nearly 900 GPCR paralogues in the human genome [5]. Their diversity facilitates responses to a large variety of external messengers such as proteins and peptides, small molecules, ions, and photons. GPCRs mediate many different cellular and physiological processes, including development, cell growth, migration, and neurotransmission [6]. Many diseases and pathological syndromes, such as hypertension, stroke, cancer, and neonatal hyperparathyroidism, are linked to GPCR inactivation, over-activation, or overexpression [7].

Structurally, previously characterized GPCRs have a single transmembrane domain, known as the GPCR domain, with seven transmembrane helices arranged as an up-down oval-shaped serpentine helix bundle with an extracellular *N*-terminus and a cytoplasmic *C*-terminus [8]. Some GPCRs also have a large extra-membrane domain, but so far none have been identified that include more than one GPCR domain. The GPCR domain relays signals from outside the membrane into the cell by undergoing conformational changes that alter interaction with cytoplasmic guanine nucleotide-binding proteins, aka, G-proteins [8]. The conformational changes are mediated by molecular switches and locks formed by several evolutionarily conserved sequence motifs, such as DRY and NPxxY [9, 10]. To induce a cellular response, the activated GPCR transduces a signal by activating its cognate G protein, which in turn activates downstream effectors, such as enzymes (adenylyl cyclase and phospholipase C-β) and ion channels. Many GPCRs form homo- or hetero-dimers, and in some cases the dimeric interaction contributes to proper signaling. For example, in class A GPCRs, the largest and most diverse of the six classes of GPCRS [11, 12], dimerization is used for regulation, whereas in class C GPCRs, which are mGlu and GABA_B_ receptors, dimerization is mandatory for activity [13-15].

If dimerization is necessary for function of certain GPCRs, there may be proteins with two or more GPCR domains concatenated on the same chain. This possibility is supported by our recent discovery of another class of multi-domain membrane proteins – bacterial outer membrane proteins with multiple beta-barrel domains [16]. To discover these bacterial multi-domain proteins, we modeled all structurally characterized outer membrane beta-barrels as hidden Markov models (HMMs) and searched in the UniRef database for proteins with non-overlapping matches to two or more beta-barrel HMMs. Here, we use a similar computational pipeline to search for proteins with multiple GPCR domains, starting with all structurally characterized GPCR domains in the ECOD database.

We discovered 57 proteins with more than one GPCR domain: 50 with two GPCR domains, six with three GPCR domains, and one with four consecutive GPCR domains linked to each other in the same protein chain. We predicted the three-dimensional structures of these proteins, which we refer to as multi-GPCRs, with AlphaFold, a machine learning tool that predicts structures at near-experimental accuracy [17]. AlphaFold uses deep networks to mine and integrate signals from internal correlations between amino acids positions of the query protein and its homologues, models the three-dimensional structure, and provides local and global estimates of the model accuracy. AlphaFold confidently predicted the structures of the putative multi-GPCRs, and the predicted structures are consistent with our hypothesis that these proteins have multiple transmembrane domains based on homology to single GPCR domains. These putative multi-GPCRs are found in various eukaryotes including fish, amphibians, birds, and mammals, specifically rodents, camels, and new world monkeys.

## Results

### Identification of putative multi-GPCRs

Based on preliminary examination of UniProt, we estimated that there are many proteins with multi-GPCR domains. Because such proteins have not been documented to date, our goal was not to detect all candidates with multi-GPCR domains but rather to focus on a smaller collection of proteins that could be convincingly shown to be multi-GPCRs. We designed our search strategy accordingly. We searched for long hits that matched two or more HMMs built from ECOD-classified GPCR domains (X-group 5001) in the protein sequence database UniRef90, which was built to cluster sequences to hide redundancies and includes over 130 million sequences. Of the 262,168 UniRef90 matches to any one of the GPCR HMMs, only matches to 2,240 proteins spanned sufficiently long segments to accommodate multiple GPCR domains. We clustered these based on their sequence similarity with CD-HIT [18]. To reduce the risk of including erroneous sequences (e.g., due to sequencing errors), we kept only proteins whose variants were found multiple times. Thus, we included only the proteins in the 51 clusters with more than five members each: 463 proteins in total. Because CD-HIT produces clusters that may be heterogeneous in terms of sequence lengths [18], which would group together proteins with different numbers of GPCR domains, we further split each of the clusters to sub-clusters of similar sequence lengths.

We applied a final filter to these clusters: We plotted the number of aligned ECOD GPCRs for each residue as a function of its position in the sequence (Figure S1). Then, we visually inspected the resulting plot to verify that there were multiple stretches, each longer than 200 residues, where the residues aligned to many overlapping GPCRs. Sixteen clusters satisfied this requirement (Figure S1). The sequences that did not satisfy this requirement were excluded. As an example, in one of the sequences excluded, there were two segments along its sequence that aligned with GPCR HMMs. However, only one has about 200 residues, whereas the other, with only about 100 residues, is too short to accommodate a full GPCR domain (Figure S2). Table 1 lists the 57 proteins, in 16 clusters, that passed all these tests. Interestingly, some of their single-GPCR domain homologues are known to form dimers (Table S1), indicating that the consecutive GPCR domains within a multi-GPCR protein may physically interact.

**Table 1:**
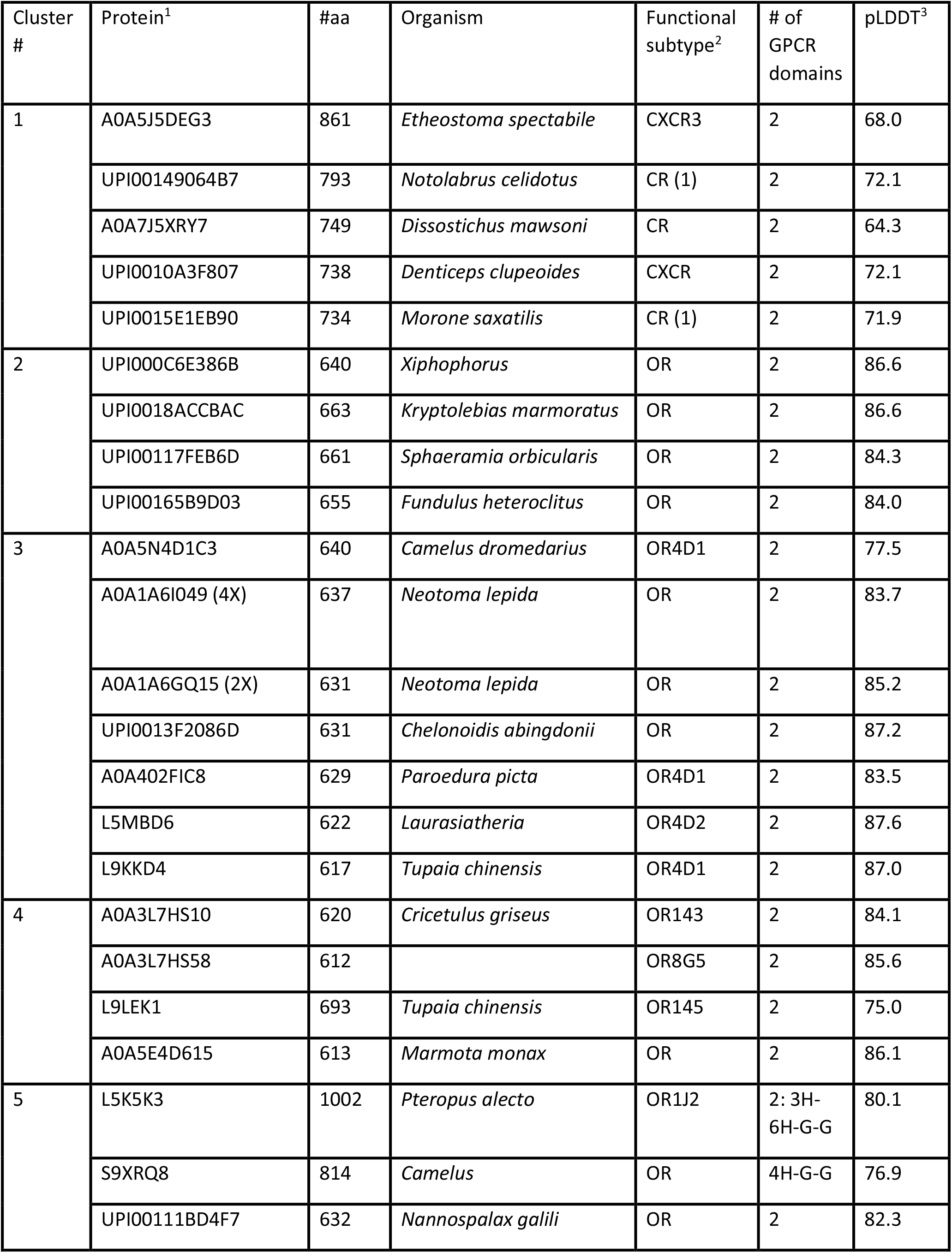
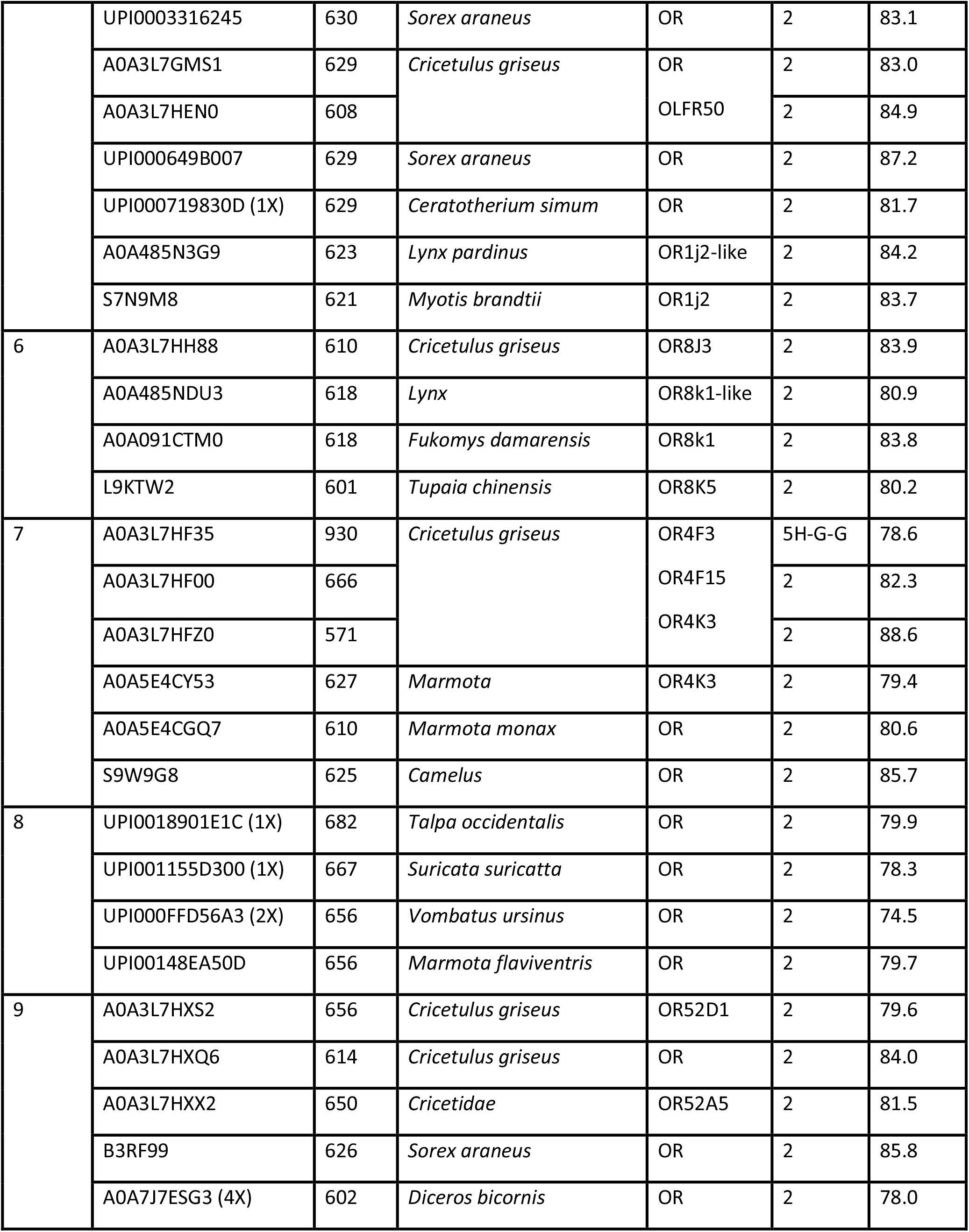
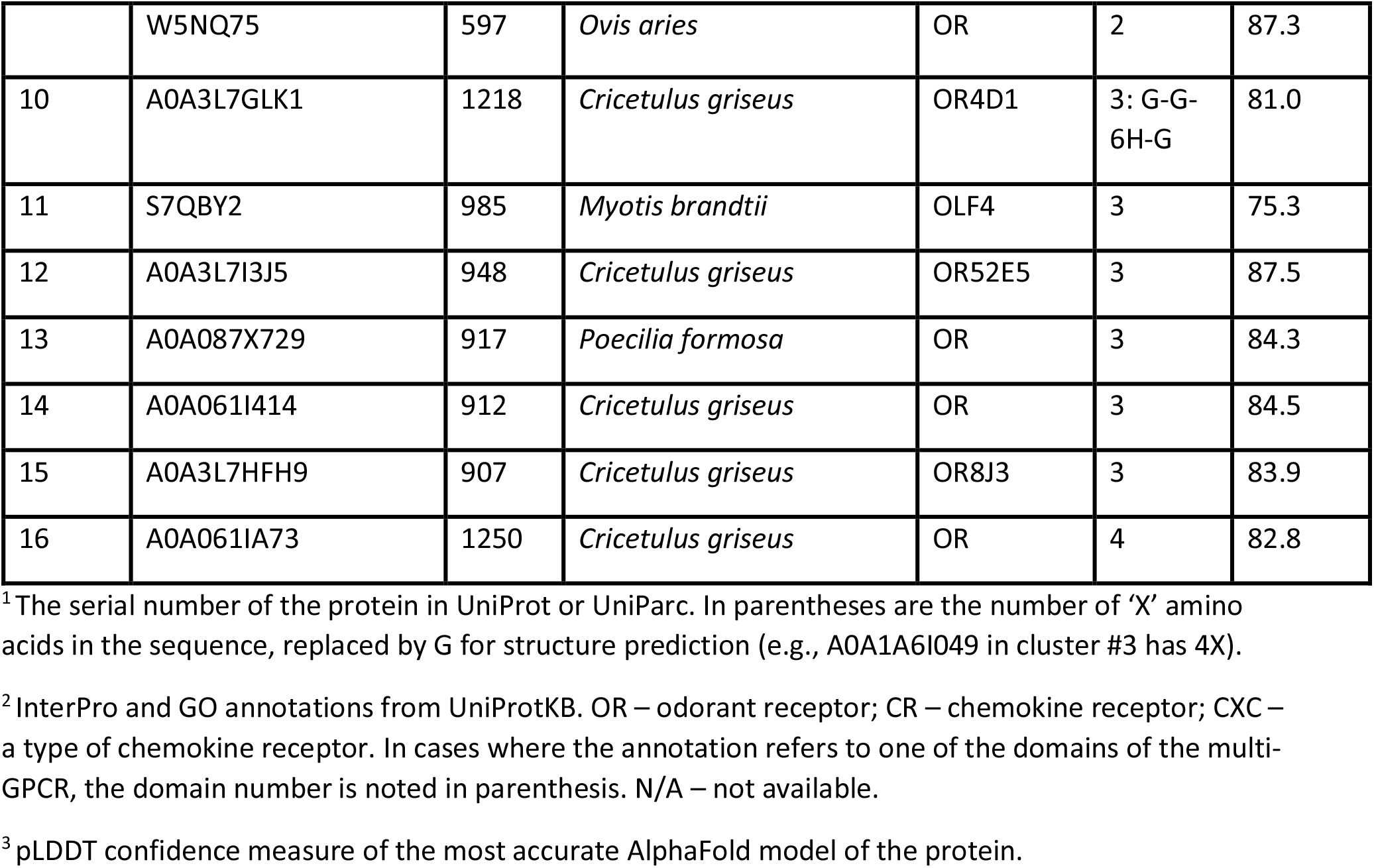
The 57 multi-GPCRs.

### AlphaFold predictions of structures of multi-GPCRs

Using AlphaFold [17], we predicted the structures of the identified 57 putative multi-GPCR proteins. In the seven proteins where an unknown amino acid (X) was listed, we replaced it with a glycine (G) for the prediction. We ran AlphaFold using the predicted template modeling score (pTM), as these reliably give the predicted local-distance difference test (pLDDT) values and the predicted pairwise aligned errors. The model with the highest pLDDT score was selected for each protein. The model structures are predicted to be accurate (i.e., high pLDDT) overall: For 55 of the 57 proteins, the pLDDT score is above 70 (very good), and in 40 it is even above 80; the average is 81.6 (Table 1). Figure 1 presents colormaps of the predicted pairwise aligned errors, showing that there is high confidence in blocks of approximately 300 residues that include the different GPCR domains along the chain, but low confidence in their positioning relative to one another. When the 57 predicted structures were inspected in a molecular viewer, we observed 50 proteins with two predicted GPCR domains (clusters #1-#9), six with three predicted GPCR domains (clusters #10-#15), and one with four predicted GPCR domains (cluster #16). There are four proteins that contain GPCR-like segments that were not marked as full domains because they have fewer than seven helices: In cluster #5, the protein L5K5K3 has three helices and six helices upstream of its two GPCR domains, and protein S9XRQ8 has four helices upstream of its two GPCR domains; in cluster #7, protein A0A3L7HF35 has five helices upstream of its two GPCR domains; and in cluster #10, protein A0A3L7GLK1 has six helices between the second and third GPCR domains. Focusing only on domains that have the full-fledged seven helices, there are a total of 122 GPCR domains in the 57 proteins.

**Figure 1:**
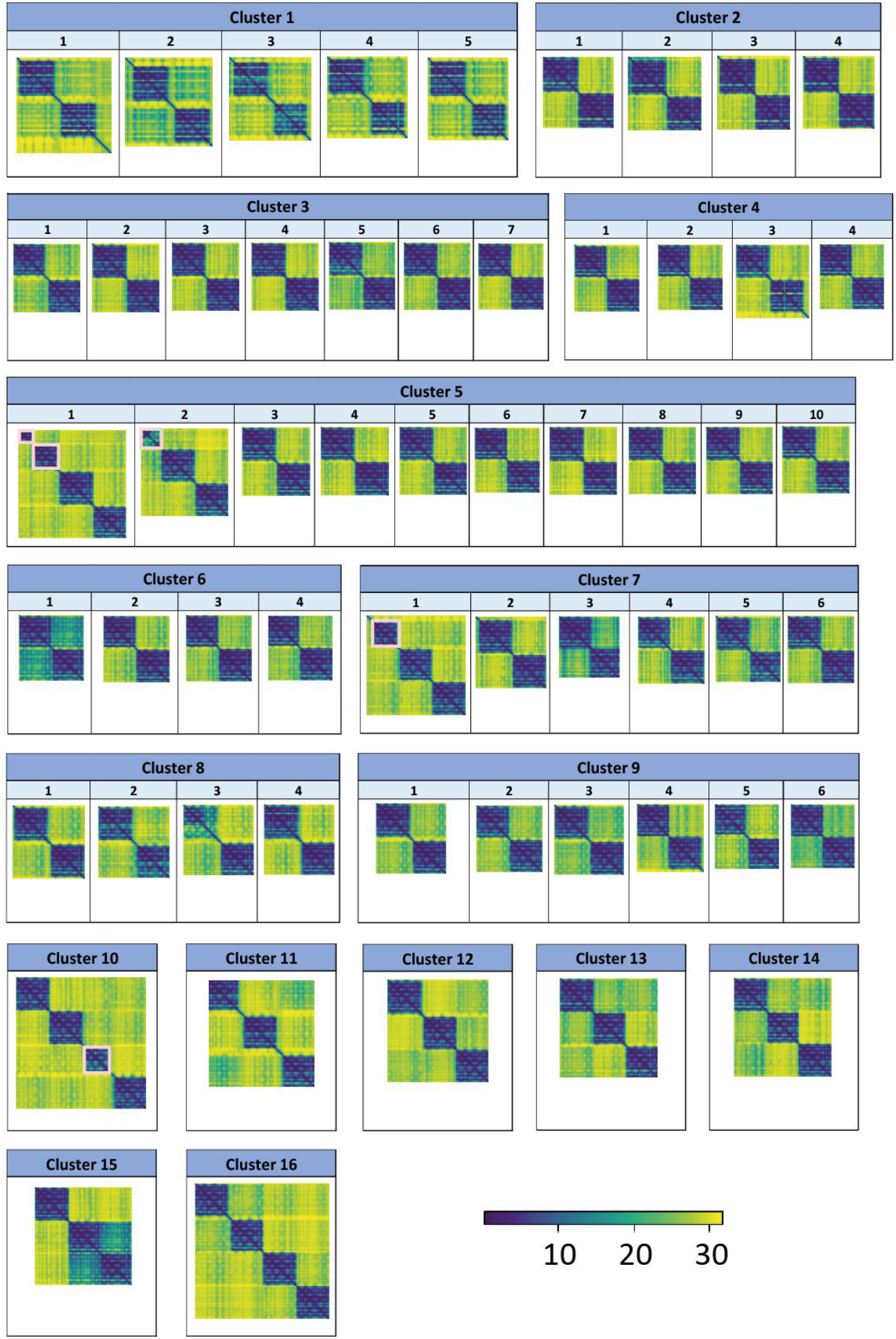
AlphaFold predicted aligned errors for the 57 multi-GPCR proteins. The estimated accuracy of the relative positions of the predicted structures are shown as colormaps of the *n*×*n* matrix of predicted aligned errors. For each protein of *n* residues, the color at (*x, y*) indicates AlphaFold’s expected position error at residue *x* if the predicted and true structures were aligned on residue *y*. AlphaFold assigns high confidence to relative amino acid positions within the individual GPCR domains (blue shades), and low confidence to the relative orientation of the domains with respect to each other (yellow shades). The colormaps are grouped by their clusters. The pink rectangles mark the four cases (first two multi-GPCRs in cluster #5, first multi-GPCR in cluster #7, and first multi-GPCR in cluster #10) with segments that have fewer than seven helices.

For each of the 57 proteins in the 16 clusters, Table 1 lists the number of amino acids, the number of GPCR domains, the organism, and the AlphaFold pTM score for the best predicted model. The PDB-format files of the best AlphaFold predicted models are available online^1^. We identified the linkers between the GPCRs; these are listed in Table S2. The average linker length is 34 amino acids. We also calculated the (average) internal similarities among the GPCRs in the same chain and verified that GPCRs in the same chain are not identical or nearly identical, which would indicated a recent duplication or sequencing error (Figure S3). The averages of sequence identity and sequence similarity are 57% and 66%, respectively.

### Phylogenetic profiles of multi-GPCRs

For each of the 57 multi-GPCR proteins, we searched for close homologues in UniRef90 to extract the list of species in which they appear. Figure 2 shows a matrix of the occurrences of the 57 proteins and the species in which we found their close homologs. Apart from one case (A0A3L7GLK1 in cluster #10 with three GPCR domains), all proteins have homologues in at least two Chordata species. Most multi-GPCRs appear in mammals, birds, reptiles, and amphibians. A minority were detected in fish. Receptors with two GPCR domains of cluster #1 are, almost without exception, exclusive to fish. The two-GPCR-domain proteins in cluster #2 appears in all families: mammals, birds, reptiles, amphibians, and fish. The rest of the receptors with two GPCR domains are generally not found in fish. We found homologues of the proteins with three GPCRs and with four GPCRs (clusters #10-16) in very few species. The number of species in which a multi-GPCR is identified decreases with sequence length (Figure S4). The most commonly found multi-GPCR protein with three predicted GPCR domains was detected in 17 species, whereas the most commonly found multi-GPCR with two predicted GPCR domains was detected in 66 species.

**Figure 2:**
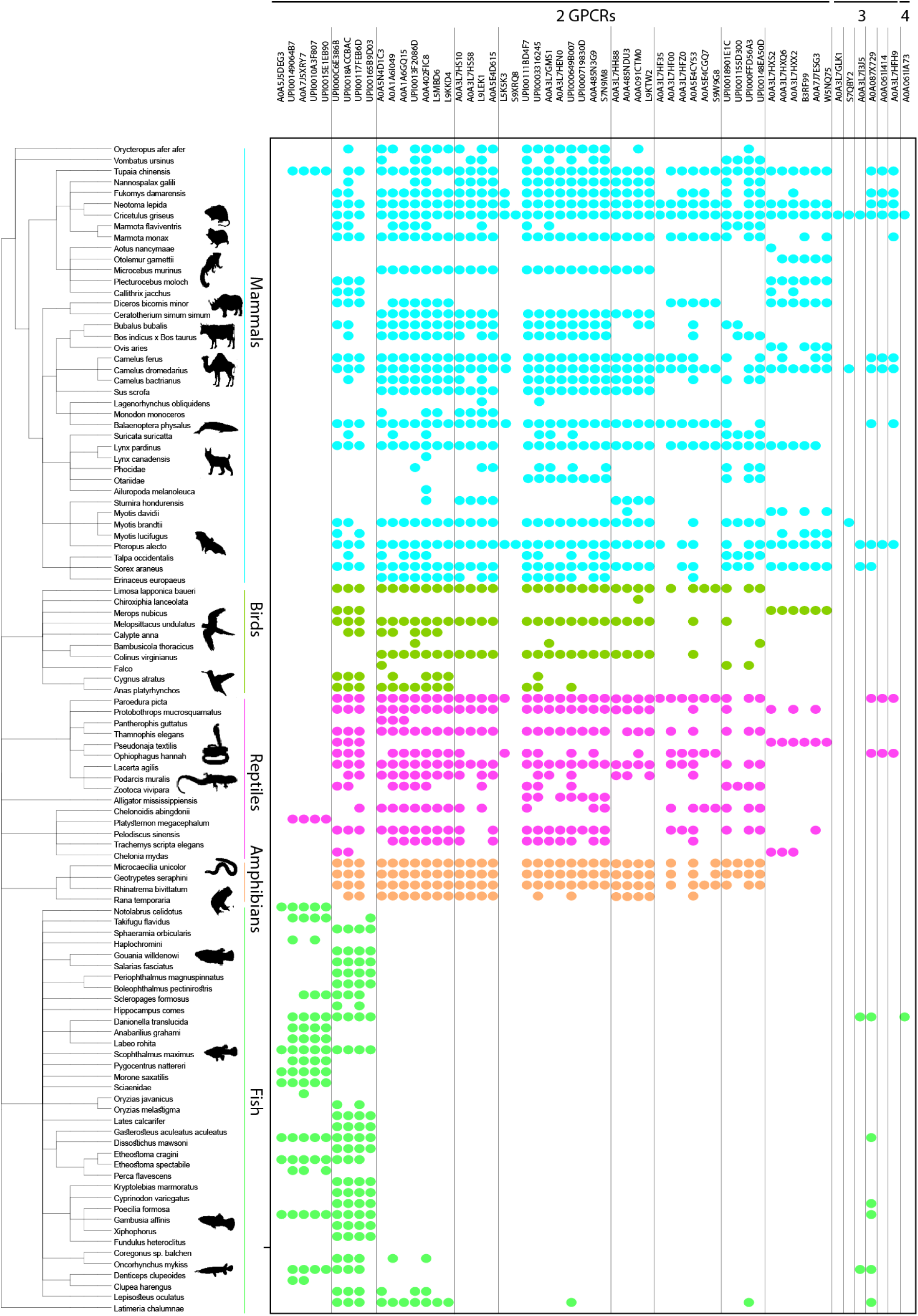
Tree of life distribution of homologues of the 57 multi-GPCR proteins. The proteins are plotted along the x-axis (first two-GPCR domains, then three-GPCR domains, then the four-GPCR domains), ordered as in Table 1. Vertical lines indicate cluster boundaries. Homologues were identified in 106 species, and are shown as leaves of a phylogenetic tree along the y-axis. Colored dots mark the species found with a homologue of a protein: cyan for mammals, green for birds, pink for reptiles, orange for amphibians, and light green for fish.

### The multi-GPCRs found are olfactory- and chemokine-related

The domains detected in multi-GPCR proteins are only homologous to domains present in muscarinic acetylcholine receptor M3 (5001.1.1.1), adenosine receptor (5001.1.1.22), rhodopsin (5001.1.1.32), and orexin receptor type 2/ chemokine receptor type 4 (5001.1.1.33) of all domains in ECOD. This holds true even though our search started with HMMs of the GPCR domains of all solved structures in ECOD. In particular, even though their HMMs were included in our search, we did not find any multi-GPCRs with homology to the glucagon receptor (5001.1.1.5), metabotropic glutamate receptor 5 (5001.1.1.6), rhodopsin (5001.1.1.7), cytochrome b562 (5001.1.1.8), or progestin and adipoQ receptors (HlyIII) (5001.1.1.12).

Table 1 lists the functional receptor type and subtype, as annotated by the InterPro database [19] or the Gene Ontology (GO) resource [20], for each of the multi-GPCRs. Based on these, almost all the multi-GPCR proteins for which we found a functional annotation are olfactory receptors. The only exceptions are the chemokine receptors of cluster #1, which are almost entirely restricted to fish. Chemokine receptors are involved in the immune response in all vertebrates [21] and have regulatory roles in fish immune responses and development [22, 23].

To examine signaling function one has to know what the ligand is. Thus, we conducted literature survey of known ligands of single-domain olfactory receptors that are homologous to some of the receptors with multi-GPCR domains (Table S3). The homology suggests that perhaps the multi-GPCR receptors respond to the same (or chemically similar) ligands.

### Conservation of functional motifs indicative of signaling activity in the multi-GPCR proteins

DRY and NPxxY are evolutionarily conserved GPCR motifs, known to participate directly in receptor activation [9, 10]. The arginine in DRY forms a salt bridge, also called an ‘ionic lock’, that stabilizes the inactive state and breaks upon receptor activation. Most of the multi-GPCR proteins that we identified have both the conserved DRY (Table S4) and NPxxY motifs (Table S5) or their close variants. Of the 122 GPCR domains in the 57 proteins, 100 have the DRY motif, ten have the DRF variant, four have the DRL variant and three more have XRY, ERY, or DRC variants. The five remaining cases are only found in four multi-GPCR proteins: the second GPCR in A0A7J5XRY7 (cluster #1), the second GPCR in UPI0015E1EB90 (cluster #1), both GPCRs in A0A5N4D1C3 (cluster #3), and the first GPCR in UPI000FFD56A3 (cluster #8), which contain the motifs QRF, DCY, DHL, DWY, and DHF, respectively. Thus, 53 of the 57 multi-GPCR proteins have either the DRY motif or a close variant thereof in all their GPCR domains. The NPxxY motif is found in 115 of the GPCR domains in the 57 proteins. In the chemokine proteins of cluster #1 there are three cases where the second GPCR has an RPxxY motif and one with an RPxxC motif. In proteins from cluster #3, the first GPCR of UPI0013F2086D has the motif NAxxY, and the second GPCR of L9KKD4 has the sequence NPxxS. In the abovementioned protein from cluster #8, UPI000FFD56A3, the first GPCR is SPxxY. The presence of both motifs in almost all the GPCR domains strongly supports our hypothesis that all these domains have signaling activity.

### Lengths and hydrophobicities of inter-GPCR linker segments

The GPCR domain has an uneven number of transmembrane helices, and for proper signaling function, it is oriented with its *N*-terminus in the outer side of the cytoplasmic membrane and its *C*-terminus in the cytoplasm. Thus, the linker between two consecutive and similarly oriented GPCR domains in a multi-GPCR protein should span the membrane to connect the cytoplasmic *C*-terminus of one domain to the extracellular *N*-terminus of the next domain. For this, the linker must be sufficiently long and hydrophobic. Not all the linkers in the multi-GPCRs we found are sufficiently long to cross the membrane: In six cases of the 57 identified proteins, the linkers are less than 15 residues (Table S2, highlighted in red). Furthermore, to cross the membrane, the linkers must be sufficiently hydrophobic. We calculated the hydrophobicity of a sliding window of 15 consecutive residues within the linkers using the Kessel & Ben-Tal scale [24] and identified the most hydrophobic window in each linker (Table S6). The Kessel & Ben-Tal scale estimates the water-to-membrane transfer free energy, with negative values favoring membrane partitioning. Surprisingly, even the most hydrophobic window within almost all linkers was too polar to cross the membrane, with calculated values larger than 1 kcal/mol and an average of 16.9 kcal/mol (Table S6). For comparison, the most hydrophobic 15-residue segment in the whole protein (including the GPCR domains) is -18.0 kcal/mol averaged over the whole dataset.

The one case with a sufficiently hydrophobic linker is A0A3L7HXX2, a protein of the Cricetidae rodent family with two GPCR domains that has homologues in 21 species. The linker in this protein has a hydrophobic segment with Kessel & Ben-Tal scale score of -4.2 kcal/mol (positions 342 through 356) (Figure 3A). The pairwise errors in the AlphaFold prediction reveal considerably more confidence in the structures of the two GPCR domains than their relative positioning or of the structure of their linker (Figure 3B). Thus, we modeled the linker manually (Figure 3C, D, E): The first 20 residues, which contain many polar and charged amino acids, are organized as a random coil, mostly outside the membrane. The remaining 25 amino acids, which are mostly hydrophobic, span the membrane in an ideal alpha-helix conformation.

**Figure 3:**
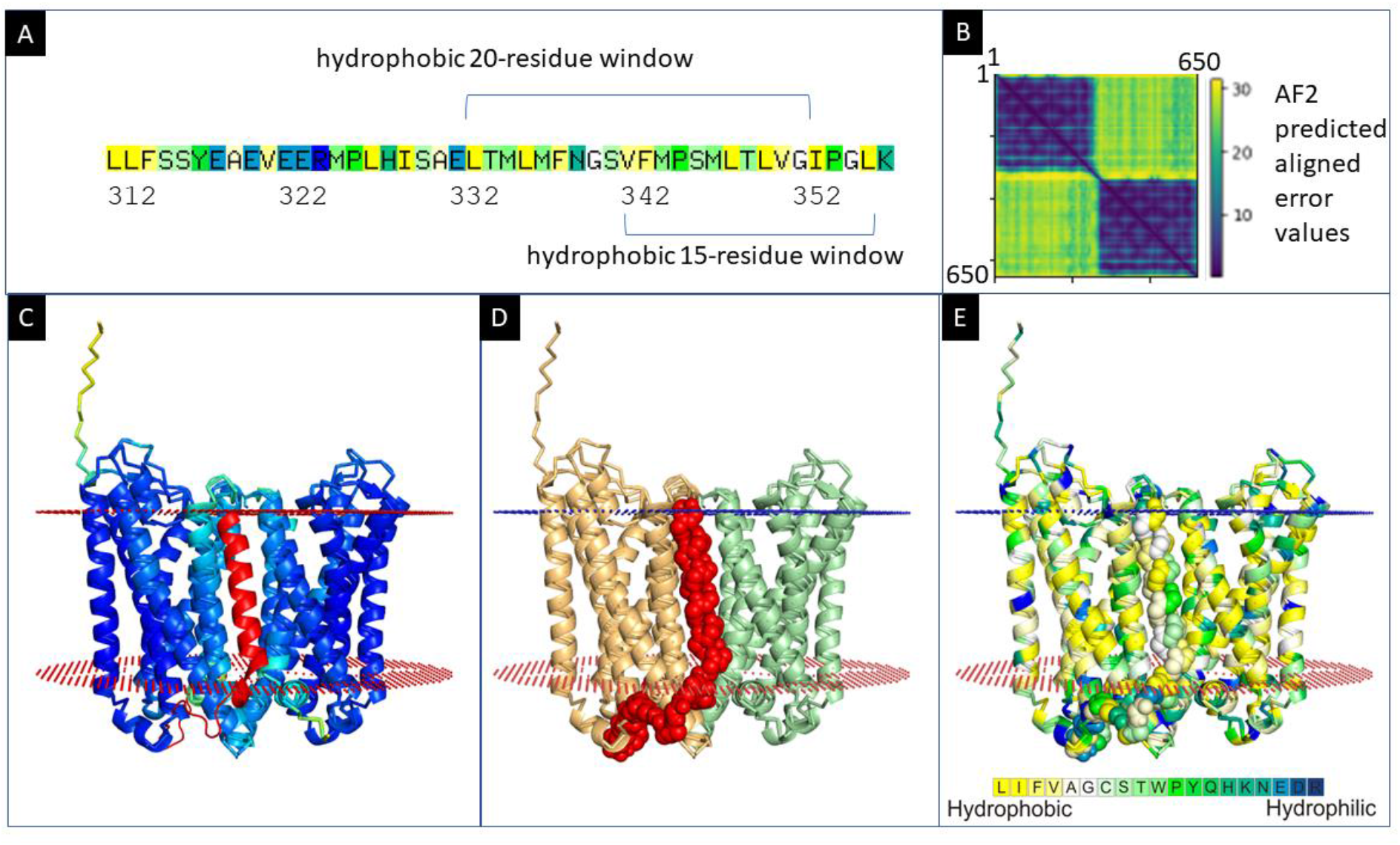
Predicted structure of two-GPCR-domain A0A3L7HXX2. (A) Kessel & Ben-Tal scale hydrophobicity scores of the linker region. The most hydrophobic 20-residue and 15-residue windows within the linker are indicated. (B) Heatmap visualization of the predicted errors in the pairwise positioning of the residues in A0A3L7HXX2 in the structure predicted by AlphaFold. The positioning of residues within each of the GPCR domains is predicted to be accurate, whereas the relative positioning of the two domains and the linker structure are not. (C) The most confident model (pTM 81.5) from AlphaFold, colored by the pLDDT confidence measure with the linker, residues 312-358 (red), modeled manually. The first 20 positions of the linker, which contain many polar and charged amino acids, are organized as a random coil mostly outside the membrane, and the remaining mostly hydrophobic 25 amino acids form an alpha-helix that crosses the membrane, placing both GPCRs in the same orientation. The extracellular (small blue spheres) and intracellular (small red spheres) boundaries of the membrane are also shown (predicted by the OPM web server). (D) Model as in panel C with the two GPCR domains colored wheat and pale green and the linker region colored red with atoms shown as spheres. (E) Model as in panel C with the structure colored by the Kessel & Ben-Tal hydrophobicity scale scores.

### Case study: the conservation of functional residues in both GPCR domains of S7QBY2

As a case study, we analyzed in detail the sequence and AlphaFold-predicted structure of S7QBY2, a protein from the bat *Myotis brandtii*. This protein of 985 amino acids has three GPCR domains. S7QBY2 was automatically annotated as an olfactory receptor in InterPro [19]. The three GPCR domains are homologous to the single GPCR domain of a known human olfactory receptor, OR7A10, with 81%, 80%, and 76% sequence identity, respectively. Sequence motifs that are associated with functionally active olfactory receptors are present and conserved in all three domains of S7QBY2, supporting the hypothesis that each of the three domains have signaling activity. Were these domains not functional, we would expect that random drift would have considerably decreased the sequence similarity. In all three GPCR domains of S7QBY2, the most significantly conserved motifs for GPCR activation [9, 10] are 3.49-D/ERY-3.51 and 7.49-NPxxY-7.53 (marked by the Ballesteros–Weinstein numbering scheme [25], with the first digit representing the number of its transmembrane helix; Tables S4 and S5). All three domains include other known olfactory receptor-associated motifs as well [26, 27]: The 6.38-FYG-6.40 motif, which resides just below the ligand binding pocket. This motif is thought to participate in the molecular mechanism that senses agonist binding. Additional motifs are3.52-VAICxPLxY-3.60 and 6.24-KAFSTCASH-6.32, which are located near the G protein-binding site of known GPCRs and which may participate in the transfer of signal from the GPCR to the G protein [26, 28]. Three additional olfactory receptor-associated motifs are also conserved in all three domains: 2.34-LHxPMY-2.39, 2.46-LSxxD-2.50, and 3.46-MAY-3.48.

All five model structures of S7QBY2 predicted by AlphaFold are very similar to each other and we therefore describe only one. In the model structure, the three GPCR domains are oriented in the same direction, with the *N*-termini facing the extracellular environment and the *C*-termini facing the cytoplasm (Figure 4A). The positively charged residues lysine and arginine are asymmetrically distributed in all three domains and are located mostly in the cytoplasmic loops, in agreement with the so-called positive-inside rule [29]. This distribution allows both sides of each domain to interact favorably with the head-groups of the lipid bilayer. The inferred G protein-binding cavities are located on the intracellular sides of each of the three domains (Figure 4B), which would allow these domains to transduce the external signal to their respective cytoplasmic G-proteins. It is noteworthy, though, that the model-structure is somewhere in between the active and inactive conformations so that it is impossible to reveal the exact details of the interaction with the G protein. The VAICxPLxY and KAFSTCASH motifs are not only conserved, but also face the predicted locations of the α5 helix of the G protein in all three domains. The predicted parallel orientations of the three domains are made possible by the two long linkers. For the linkers to be sufficiently hydrophobic to cross the membrane, we had expected that they would contain segments of at least 15 hydrophobic residues; however, this is not the case (Table S6). This structure suggests that each pair of adjacent domains would most likely be able to bind only a single G protein molecule at a time, due to the large size of the latter. Thus, our model suggests that this three-domain GPCR could be expected to bind at most two G proteins at one time.

**Figure 4:**
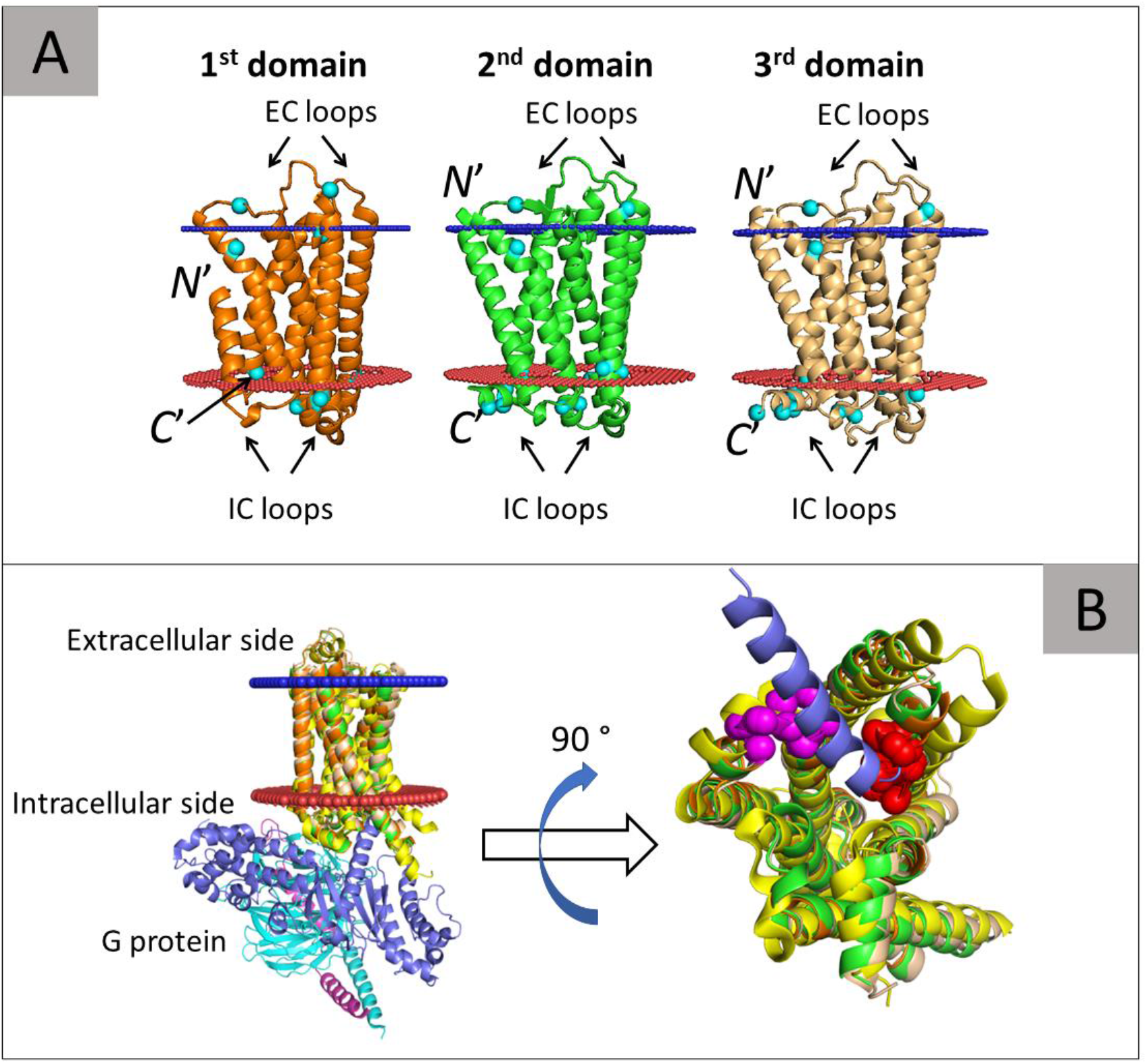
Predicted structure of three-GPCR-domain S7QBY2. (A) Predicted model with the first, second, and third successive GPCR domains of the protein colored orange, green, and wheat, respectively. The inferred extracellular and intracellular membrane boundaries are also shown (calculated by the PPM web server[35]), as blue and red spheres, respectively. The membrane topologies of the domains are shown by noting their *N*- and *C*-termini and the intracellular (IC) and extracellular (CE) loops. The arginine and lysine residues (cyan spheres) are asymmetrically distributed in keeping with the positive-inside rule [29]. (B) Left: Side view of the structure with the three GPCR domains superimposed onto the structure of the G protein-bound (activated) β2-adrenergic receptor (β_2_-AR, PDB ID: 3sn6). The β_2_-AR receptor is colored yellow and the α, β, and γ subunits of the G protein are colored blue, cyan, and magenta. Right: An enlarged view of the G protein-binding site. The β_2_-AR and superimposed domains of S7QBY2 are rotated 90° to show the binding site. For clarity, only the α5 helix of the G protein is shown; this region directly interacts with the GPCR. The image clearly shows that all three domains of S7QBY2 contain a cavity that corresponds to the G protein-binding site of the β_2_-AR. Moreover, all domains contain the VAICxPLxY and KAFSTCASH motifs (pink and red spheres, respectively), which directly interact with the α5 helix of the G protein.

### Multi-GPCRs in primates

We were particularly interested in multi-GPCRs in primates and tailored a somewhat more lax search to this end. Using this search strategy, we identified five proteins predicted to be multi-GPCRs (Figure S5): B0VXB0 in *Callithrix jacchus*, the white-tufted-ear marmoset, and B1MT73, B1MT94, and B1MT76 in *Plecturocebus moloch*, the dusky titi monkey, and A0A1D5QNK2 in *Macaca mulatta*, the rhesus monkey.

## Discussion

Our work expands our knowledge regarding the repertoire of GPCRs. Previously, only GPCRs with a single GPCR domain per chain, occasionally including also water-soluble domains, but not with additional and autonomous transmembrane domains, had been described. We reasoned that since many GPCRs form functional dimers [13-15], there might be protein chains with multiple GPCR domains. We discovered 57 proteins from Chordata species that contain two, three, or four GPCR domains. We offer evidence that the multi-GPCRs we identified do indeed contain more than one functional GPCR domain. First, we found several homologues for each of the proteins identified. Second, the proteins were found in multiple species. Third, high-confidence AlphaFold structure predictions are indicative of expected GPCR domain folds. Furthermore, important function-related residues in the GPCR domains of the multi-GPCR proteins are conserved. That sequence motifs known to be critical for signaling activity function of receptors with single GPCR domains are conserved, suggests that the multiple GPCR domains in the proteins we identified as multi-GPCRs may also have signal transduction function. Finally, our homology-based search also detected several fragmented GPCR domains that may be involved in signal regulation, especially when present next to an intact GPCR domain.

The multi-GPCRs that we found appear only in mammals, birds, reptiles, amphibians, and fish, whereas single-domain GPCRs have much broader distribution in eukaryotes, from primitive yeast and choanoflagellate species to animals. Almost all multi-GPCRs have two GPCRs, a few have three, and one has four GPCR domains. Based on the functions of the single-domain GPCRs that are homologous to the domains we identified in the multi-GPCRs, most of the multi-GPCRs have olfactory-related function despite searching with single-domain GPCRs that have other functions as well. Olfactory-related motifs are also conserved in the identified multi-GPCRs. Linkage of multiple olfactory domains may contribute to the ability of the protein to detect and respond to various agonists, explaining the superiority of the sense of smell in some species. Evolution may have taken advantage of the diversity of GPCR domains to allow combinatorial ligand recognition within the same chain. The functions of the multi-GPCRs in one cluster (cluster #1), which are proteins found in fish, are likely chemokine related. Chemokines mediate the response of the immune system [21-23], and one can also imagine the opportunities provided by combinatorial linkage of chemokine receptors.

We recently identified proteins from another class of membrane proteins – multiple outer membrane beta-barrels (OMBBs) – with multi-domain architectures [16]. We discovered more than 30 multi-OMBBs in Gram-negative bacteria, some with up to 11 OMBB domains in one chain. Like GPCRs, protein with OMBB domains are known to form assemblies [30], which led us to speculate that multi-OMBBs might exist. Because single domain OMBBs have an even number of beta strands, the *N*- and *C*-termini of each domain are on the same (periplasmic) side of the membrane. Thus, when multiple barrels are present within the same protein, consecutive barrels are in the same orientation with respect to the membrane, as the end of the one domain is on the same side of the membrane as the beginning of the following one.

In contrast, GPCR domains, with their seven helices, cross the membrane an odd number of times, placing their *N*- and *C*-terminals on opposite sides of the membrane. The known functional mechanism of GPCR domains necessitates that they be oriented in the membrane with the *N*-terminus facing outward and the *C*-terminus facing inward, so that they can bind their external ligands and transduce signal to their cognate G proteins in the cytoplasm [8]. Thus, if the first GPCR domain is positioned properly within the membrane, the linker between the two domains must cross the membrane so that the *N*-terminus of the next domain will also be on the outward side of the membrane. For that, the linker must be sufficiently long and sufficiently hydrophobic. Most of the putative transmembrane spanning linkers in the multi-GPCRs we identified do not meet these criteria, even though we used a window of only 15 residues to evaluate hydrophobicity, far shorter than the 20 amino acids required to span the 30 Å thickness of the hydrocarbon region of the lipid bilayer if the linker residues adopt an alpha-helix conformation [24]. Alternatively, if the linker does not cross the membrane, consecutive GPCRs in the same chain will be positioned in opposite orientations with respect to the membrane. This is in agreement with many of AlphaFold-predicted structures of the multi-GPCRs we identified, albeit confidence in these relative positionings is not high. This is puzzling, and we speculate about possible resolutions for this discrepancy, focusing on the most common case of a receptor with two consecutive GPCR domains (Figure 5).

**Figure 5:**
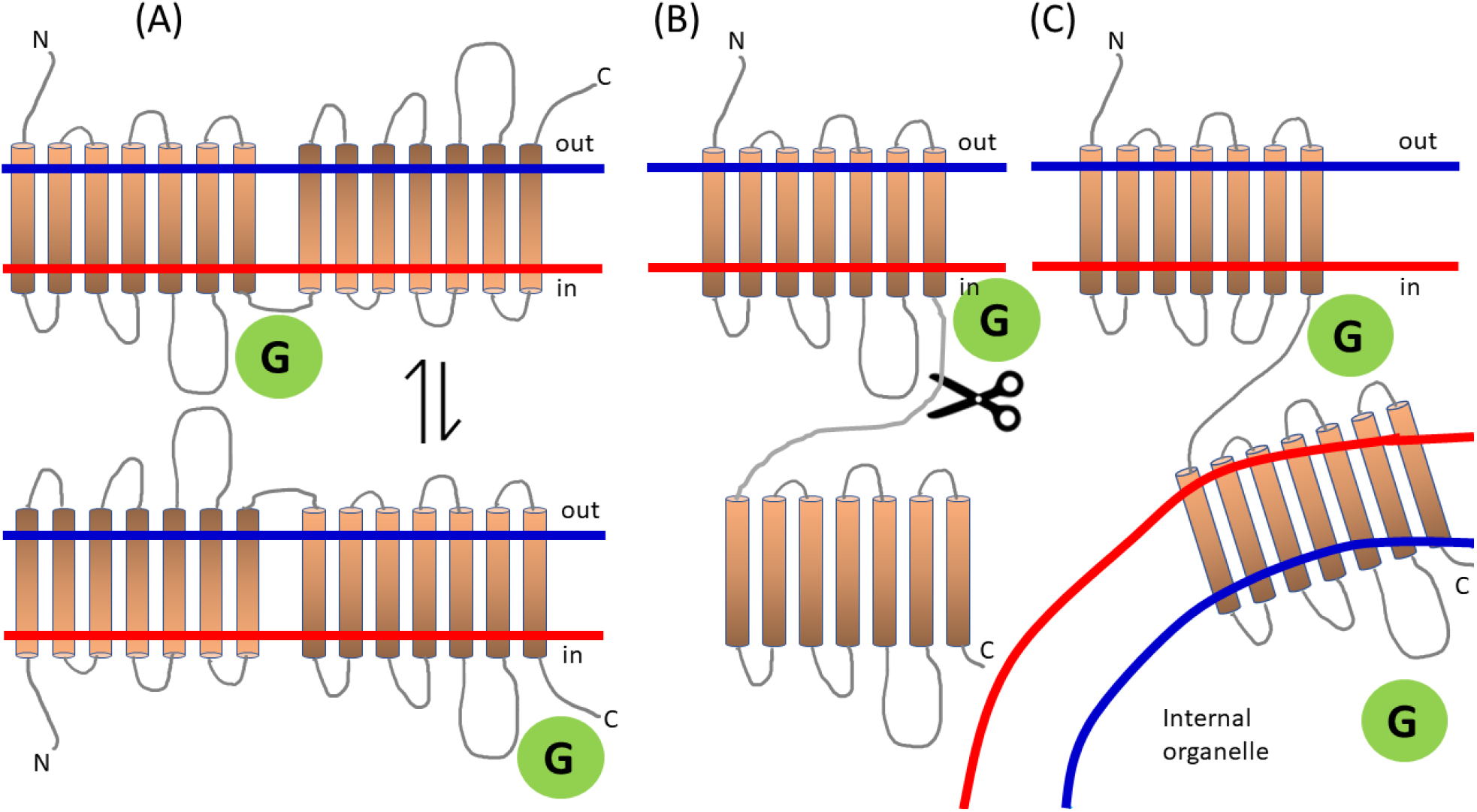
Speculative scenarios for a two-GPCR protein with a linker that is too polar to cross the membrane. (A, top) The first 7TM GPCR domain has its N terminus on the outer side, in the conventional membrane topology. We indicated the correct orientation with the color gradient. The second 7TM GPCR domain, on the other hand, is oriented in the opposite direction. Here, only the first domain is functional, and if the orientation of the protein in the membrane is reversed (A, bottom), only the second domain would be functional. (B) A (yet to be discovered) enzyme may cleave the chain. (C) The first 7TM GPCR domain may be oriented properly in the membrane, while the second is in the membrane of a nearby internal organelle.

One possibility is that the two consecutive GPCRs reside in opposite membrane topologies in the intact protein (Figure 5A). If this is the case, the number of arginine and lysine residues in the two sides of the entire protein will be similar, and the positive-inside rule would not dictate any preferred membrane topology. Thus, the protein population may partition in opposite membrane topologies, and in each protein only one of the GPCR domains will be positioned in a functionally competent membrane topology at any given time – akin to a reversible jacket. Without clear topology preference, there would be evolutionary pressure to retain the functional sequence motifs of both GPCR domains. The possible evolutionary advantages of this are that GPCR activity would be robust to membrane topology changes and that the inter-domain interaction should increase thermal stability, as is the case for protein oligomers [31].

Alternatively, the segment linking the GPCRs may be post-translationally cleaved (Figure 5B), allowing the two domains to assume the same, biologically active, orientation within the membrane. An advantage here would be that the two GPCR domains should remain in close proximity, ready to conduct joint functions. A (yet to be discovered) enzyme could carry out the cleavage, perhaps even on the membrane, and this enzyme could serve as a regulatory measure to facilitate the activation of the additional domain as an individual GPCR. Thus, the regulated cleavage of multi-GPCRs may be yet another means used by organisms to amplify specific GPCR activities in certain cells or tissues.

A third possibility is that the two GPCR domains reside in different cellular membranes, with the intact linking segment not passing through any membrane (Figure 5C). For example, one domain could partition into the plasma membrane, and the second into the membrane of an adjacent internal, vesicular organelle (e.g., a lysosome or a peroxisome) with its ligand-binding site facing the cytosol and its G protein-binding site facing the organelle’s lumen. This hypothetical process could be part of a signaling event that relocates an organelle close to the plasma membrane and then affects the internal environment of the organelle. In support of this hypothesis, GPCRs have been found in all membranous organelles, and in some of them (e.g., mitochondria, endoplasmic reticulum, and Golgi apparatus) the resident GPCRs have also been shown to exert signaling cascades into the cytoplasm or organelle lumen [32]. The latter case requires that the organelle lumen contain the other components of the signaling pathway including G proteins [32]. The overall membrane asymmetry is retained in internal organelles, where the cytoplasmic-facing side is enriched with acidic lipids [33, 34]. We speculate that the second GPCR domain would partition into the organelle membrane with its “positive side” in the organelle’s interior (i.e., lumen), which is deprived of acidic lipids. It is, however, unclear whether a “positive in the cytoplasm” rule exists for proteins in organelle membranes.

Our search pipeline had some limitations. First, to consolidate our discovery of proteins with multi-GPCR domains based on their high similarity to single-domain GPCRs of known structure, we used AlphaFold to predict structures. One may wonder how trustworthy the AlphaFold predictions are, given that the multi-domain GPCRs are very similar to single domain GPCRs of known structure. These structures are almost surely present in AlphaFold’s training data, and, given the high sequence similarity, it is possible that AlphaFold would tend to miss ways in which the GPCR domains from multi-GPCRS differ from the domains in single-domain GPCRs. This is especially likely if the multiple sequence alignments include mostly homologs of the single domains and only few homologs of multi-domain proteins. Furthermore, AlphaFold uses structural templates when available. In view of the many available structures of close single-domain homologues, the prediction likely translates, in essence, to homology modeling. Given the high sequence similarity between each of GPCR domains in our multi-GPCR proteins and single-domain GPCRs of know structure, it is hard to believe that the structures deviate from the known seven-transmembrane-helix fold. Another limitation is related to the phylogenetic profiles. We used a very strict similarity threshold to detect appearances of proteins multi-GPCR domains in the various taxa. Thus, we might have overlooked some species, and also the relative proportions of multi-GPCRs in the various species groups might be inaccurate. Obviously, experimental validation is needed to make sure that the newly discovered multi-GPCR proteins are functional. To this end, we tried to infer their ligands based on known ligands of their single-GPCR domain homologues (Table S3).

Despite these limitations, the conservation of motifs in the identified GPCR domains of the multi-GPCR proteins suggests that the multi-GPCRs we identified, e.g., in fish, camel, marmite, Chinese hamster, and new world monkeys are functional. The identification of receptors with multiple GPCR domains linked to each other in the same protein chain introduces the possibility of a novel layer of complexity in cellular signaling, providing multiple opportunities for amplification and regulation.

## Methods

We constructed HMMER profiles for all 92 GPCR domains in the 99% NR ECOD dataset (X-group 5001); we removed e2lotA1 because it has only 64 residues. We searched for each of these sequences with HHblits vs. Uniclust30 (with -cov 90 -qid 30 flags), converted the resulting alignment to Stockholm format, and built an HMM (with hmmbuild command). Then, we searched (using hmmsearch and default parameters) for each one of these 92 HMMs vs. UniRef90. We parsed the output files to remove cases for which the range of the matching residues in the GPCR query and in the UniRef90 target are both greater than 150 residues. We chose this threshold because the shortest of the 92 GPCR domains has 196 residues.

To identify protein sequences with sufficiently long matches, we counted the total number of residues in the UniRef90 target that were matched to any of the HMMs, the number of matching HMMs, the average matched length, and the average and maximal E-values. We then kept for further consideration only those where the difference between the index of the greatest matched residue (which corresponds to the end of the *C*-terminal GPCR domain) and the smallest matched residue (which correspond to the first residue of the *N*-terminal GPCR domain) was greater than 550. These amino acid segments are long enough to accommodate at least two GPCR domains separated by a linker. Then, we clustered these 2,240 sequences with CD-HIT using default parameters except that the sequence identity cutoff was 0.5. We then selected the 51 clusters with more than five sequences from the total 1395 clusters.

Less stringent search criteria were used to identify multi-GPCRs in primates. We searched against all ∼1.4 million primate sequences (taxonomy: Primates [9443]) in UniProt. We relaxed the filter to require only 500 of the residues in the sequence to align to any of the known GPCRs. We found 63 preliminary candidates. These candidates did not pass the filter of having more than five homologues, but we nonetheless predicted their AlphaFold model structures. By inspecting the model structures and the heatmaps of the predicted pairwise aligned errors, we identified five proteins predicted to be two-GPCRs (Figure S5). Of these, four were already present in our previous dataset (of sequences found when searching UniRef90) in three singleton clusters and in a cluster of three sequences. The singletons were B0VXB0 from *Callithrix jacchus*, B1MT73 from *Plecturocebus moloch*, and B1MT94 from *Plecturocebus moloch*. B1MT76 was in a cluster of three sequences that also included proteins from *Cricetulus griseus* (Chinese hamster) and *Lynx pardinus* (Iberian lynx). The fifth protein is A0A1D5QNK2 from *Macaca mulatta* (rhesus macaque).

For each protein identified using our procedure, we retrieved the receptor type and subtype using the GO and InterPro annotations in UniProtKB and UniParc.

## Acknowledgements

This research was supported by grant 1764/21 from the Israeli Science Foundation (ISF). NB-T’s research is supported in part by the Abraham E. Kazan Chair in Structural Biology, Tel Aviv University.

## Footnotes

1 https://trachel-srv.cs.haifa.ac.il/rachel/MGPCR/Table1.html

