## Supplementary data for "Proteins with multiple G protein-coupled receptor domains"

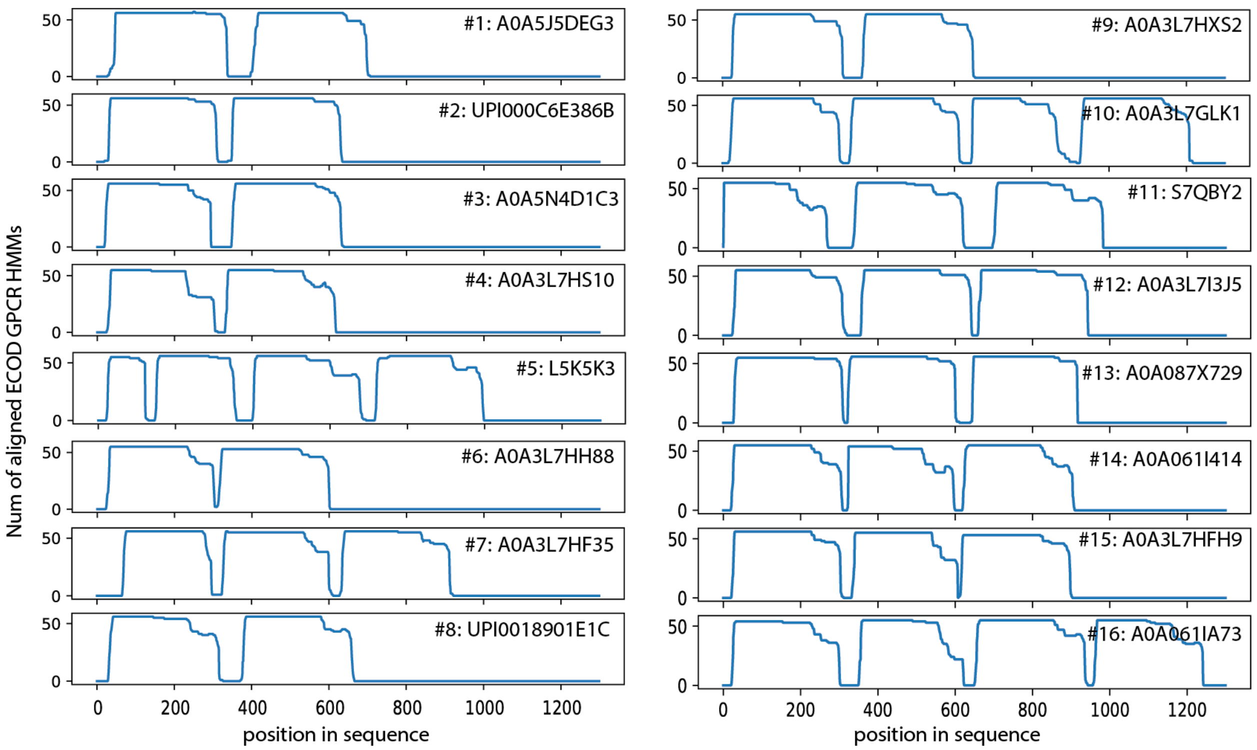


*Figure S1:* The number of ECOD GPCR HMMs aligned to each position in sixteen representatives of clusters with multi-GPCRs. In each one, we can see stretches that are longer than 200 residues with multiple matches. For each panel, the number of cluster and the representative protein is listed. In each one we see multiple stretches that are candidates for GPCR segments. Notice that in some of these proteins there are long stretches that are shorter than a full GPCR (e.g., in cluster #5), as described in the text.

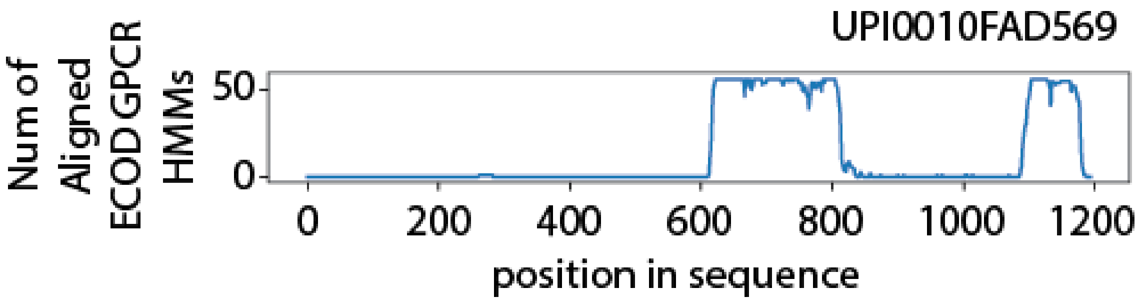


*Figure S2:* The sequence UPI0010FAD569 was filtered out because there was only a single sufficiently-long segment along the sequence that matches to the ECOD GPCR HMMs (between residues 610-837). The stretch between residues 1086-1184 is not long enough. Thus, we removed this sequence, and 33 additional (shorter) sequences that were in its CD-HIT cluster.

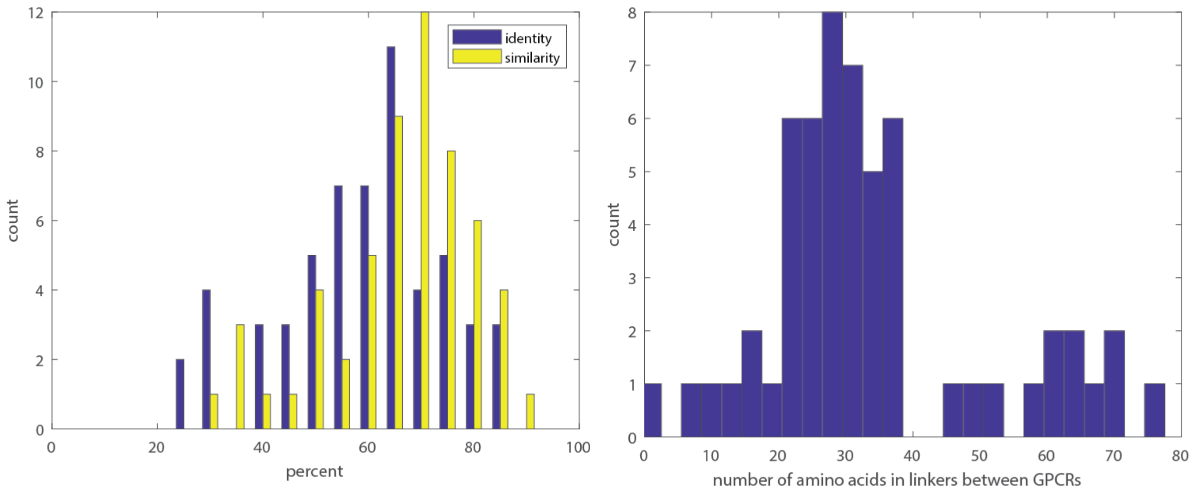


Figure S3: (A) Distribution of sequence identify/similarity among the different GPCR domains within the same chain.
(B) Distribution of the number of residues in the linker regions between the different GPCR domains in the same chain.

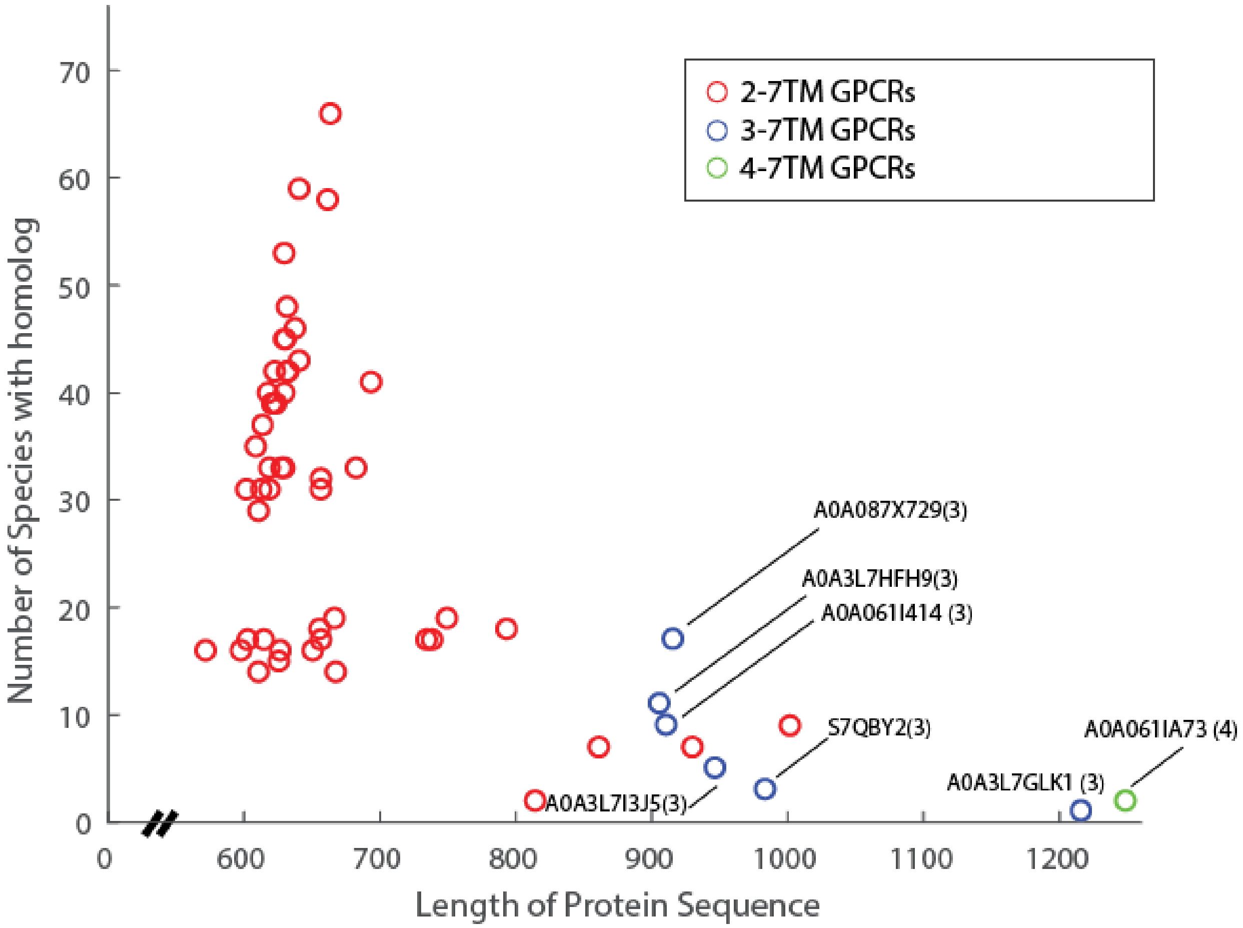


Figure S4: The number of species with a detectable multi-GPCR receptor decreases with the length of protein sequence . The number of residues in the set of 57 proteins with multi GPCR domains ranges from 571 to 1250 (x-axis), and the number of species ranges from 1 to 75 (y-axis). The identities of the 6 proteins with 3 GPCR domains, and the single protein with 4 GPCR domains are marked.

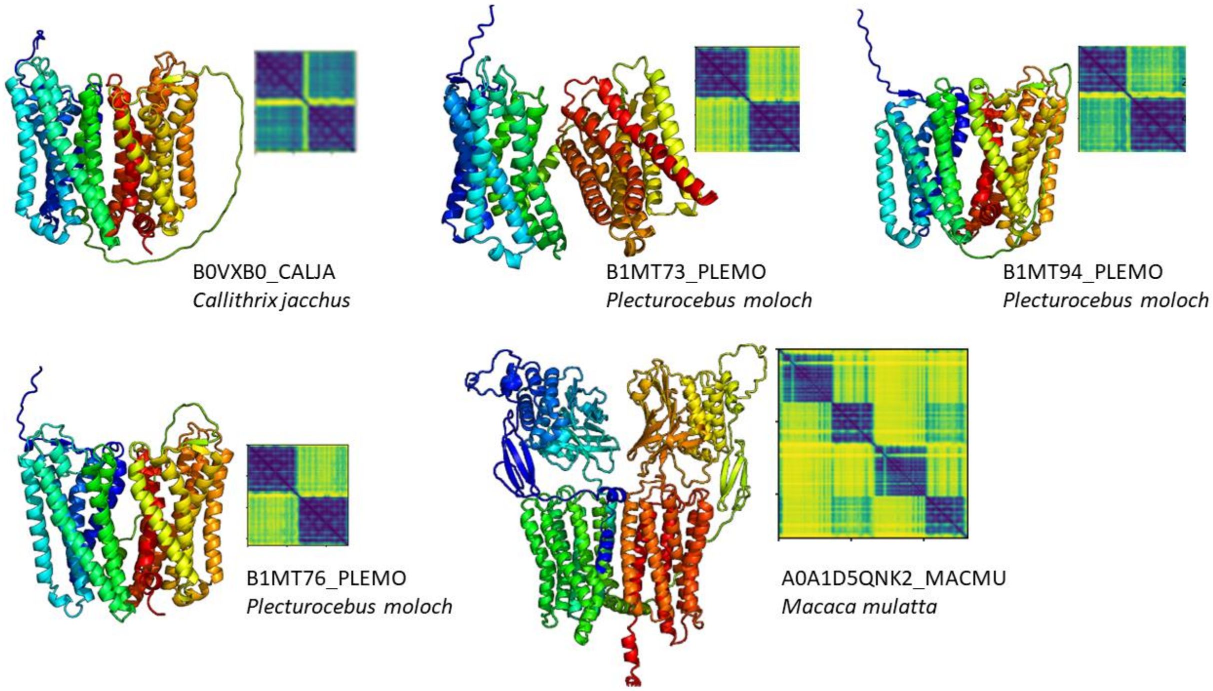


*Figure S5:* The best AlphaFold models and the predicted error for the five multi-GPCR primate proteins found

**Table S1. Single-domain GPCRs, homologous to some of the multi-GPCRs reported here, which are known to form dimers (see [3] and references therein).** The data refers to specific GPCR types that are homologous to some of the multi-GPCRs discovered in this study, and which are found in the ECOD database. The ECOD IDs that correspond to each GPCR type are specified.

| GPCR type | ECOD IDs |
|---|---|
| Rhodopsin | e4zwjB1<br>e1jfpA1<br>e1u19A1 |
| Beta-1 adrenergic receptor | e4bvnA1<br>e2vt4A1 |
| Beta-2 adrenergic receptor | e2r4rA2<br>e2rh1A4<br>e3kj6A2<br>e3p0gA1<br>e3sn6R4<br>e4ldlA3 |
| Muscarinic acetylcholine receptor | e4dajA3<br>e4u15B1 |
| Histamine H1 receptor | e3rzeA3 |
| $A_{2A}$ adenosine receptor | e2ydvA1<br>e3emlA4<br>e3reyA1<br>e3vgaA2<br>e4eiyA3 |
| Angiotensin II receptor (AT1) | e4yayA1 |
| Lysophosphatidic acid receptor | e4z34A2<br>e4z36A1 |

**Table S2: Lengths of linkers -- see attached Excel file**

**Table S3. Possible agonists for some of the multi-GPCRs, inferred from literature survey of their single GPCR homologues.** The multi-GPCRs shown here (column 1) are all olfactory receptors (ORs). For each OR type (column 2), the table specifics the name of the agonist reported for a single-domain GPCR (column 3), as well as its PubChem (column 4) and CAS (column 5) numbers. When available, the log EC50 value (column 6) is shown as well.

| 1 | 2 | 3 | 4 | 5 | 6 | 7 |
|---|---|---|---|---|---|---|
| **Multi-GPCRs** | **OR** | **Agonist** | **PubChem** | **CAS no.** | **log EC50** | **Database/Ref** |
| A0A5N4D1C3<br>A0A402FIC8<br>L9KKD4<br>A0A3L7GLK1 | OR4D1 | androstenone | 6852393 | 18339-16-7 | n.a. | primaryodors.org |
| L5MBD6 | OR4D2 | eugenol acetate | 7136 | 93-28-7 | -5.0 | [1] |
| A0A091CTM0<br>A0A485NDU3 | OR8K1 | nonanal | 31289 | 124-19-6 | -5.5 | primaryodors.org |
| L9KTW2 | OR8K5 | (R)-carvone | 439570 | 6485-40-1 | -5.3 | [2] |
| A0A3L7HFZ0<br>A0A5E4CY53 | OR4K3 | eugenol | 3314 | 97-53-0 | -3.3 | primaryodors.org |
| A0A3L7HXS2 | OR52D1 | methyl octanoate | 8091 | 111-11-5 | -1.1 | primaryodors.org |

**Table S4: The DRY motifs (padded by 4 residues on each side) -- see attached Excel file**

**Table S5: The NPxxY motifs (padded by 3 residues on each side) -- see attached Excel file**

**Table S6: Most hydrophobic 15-residue window according to the Kessel-Ben-Tal scale – see attached Excel file**
